# A population density and moment-based approach to modeling domain Ca^2+^-mediated inactivation of L-type Ca^2+^ channels

**DOI:** 10.1101/014449

**Authors:** Xiao Wang, Kiah Hardcastle, Seth H. Weinberg, Gregory D. Smith

**Affiliations:** Department of Applied Science, The College of William & Mary, Williamsburg, VA, USA

**Keywords:** L-type Ca^2+^ channel, population density model, moment-based model, Ca^2+^-dependent inactivation

## Abstract

We present a population density and moment-based description of the stochastic dynamics of domain Ca^2+^-mediated inactivation of L-type Ca^2+^ channels. Our approach accounts for the effect of heterogeneity of local Ca^2+^ signals on whole cell Ca^2+^ currents; however, in contrast with prior work, e.g., Sherman et al. (1990), we do not assume that Ca^2+^ domain formation and collapse are fast compared to channel gating. We demonstrate the population density and moment-based modeling approaches using a 12-state Markov chain model of an L-type Ca^2+^ channel introduced by Greenstein and Winslow (2002). Simulated whole cell voltage clamp responses yield an inactivation function for the whole cell Ca^2+^ current that agrees with the traditional approach when domain dynamics are fast. We analyze the voltage-dependence of Ca^2+^ inactivation that may occur via slow heterogeneous domains. Next, we find that when channel permeability is held constant, Ca^2+^-mediated inactivation of L-type channel increases as the domain time constant increases, because a slow domain collapse rate leads to increased mean domain [Ca^2+^] near open channels; conversely, when the maximum domain [Ca^2+^] is held constant, inactivation decreases as the domain time constant increases. Comparison of simulation results using population densities and moment equations confirms the computational efficiency of the moment-based approach, and enables the validation of two distinct methods of truncating and closing the open system of moment equations. In general, a slow domain time constant requires higher order moment truncation for agreement between moment-based and population density simulations.

## Introduction

Voltage-gated Ca^2+^ channels fall into three main groups: Ca_v_1 (L-type, L for “long lasting”), Ca_v_2 (P-, N-, and R-type), and Ca_v_3 (T-type, T for “transient”) [1]. Among them, plasma membrane L-type Ca^2+^ channels (LCCs) are widely expressed in many tissues and are known to play an important role in Ca^2+^-dependent responses of electrically excitable cells. In cardiac myocytes, for example, Ca^2+^ influx via L-type Ca^2+^ channels into the dyadic subspace triggers sarcoplasmic reticulum (SR) Ca^2+^ release and muscle cell contraction [2–4]. L-type Ca^2+^ channels also play a key role in coupling synaptic excitation to activation of transcriptional events that contribute to neuronal plasticity [5]. The activation of LCCs is voltage-dependent while the inactivation occurs via both voltage-and Ca^2+^-dependent mechanisms; consequently, the formation of Ca^2+^ microdomains following LCC influx can greatly influence the stochastic gating of LCCs and the physiology of excitable cells [6,7].

There are four subtypes of LCCs that are denoted Ca_v_1.1–1.4. Ca_v_ 1.1 is primarily found in skeletal muscle and Ca_v_ 1.4 is mainly found in retinal cells [8,9]. Ca_v_ 1.2 and 1.3 are highly expressed in cardiac myocytes and cells of the central nervous system [10,11]. In neuroendocrine cells, Ca_v_1.2 and 1.3 are both involved in action potential generation, bursting activity and hormone secretion [8,12]. Ca_v_1.3 is biophysically and pharmacologically distinct from Ca_v_1.2. For example, Ca_v_1.3 activates at a more hyperpolarized voltage, has faster activation, and slower and less complete voltage-dependent inactivation than Ca_v_1.2 [13,14]. In the heart, Ca_v_1.2-mediated Ca^2+^ currents play an important role in systolic events such as EC coupling (the triggered release of SR Ca^2+^) [15] and the plateau depolarization (phase 2) of the action potential [16]. Ca_v_ 1.3, on the other hand, is highly expressed in cardiac pacemaker cells and is the major regulator of RyR-dependent local Ca^2+^ release during the diastolic phase [17]. Inactivation of Ca_v_1.2 channels is both voltage-and Ca^2+^-dependent [7]; however, certain Ca_v_1.4 L-type channels do not exhibit Ca^2+^-dependent inactivation [8]. L-type Ca^2+^ channels that undergo Ca^2+^-dependent inactivation do not in fact result in long lasting currents, in spite of the traditional nomenclature [8].

Models of Ca^2+^-inactivation often assume a high density of Ca^2+^ channels and the slow accumulation of intracellular Ca^2+^ in a cortical shell near the plasma membrane [18]. In the context of low density Ca^2+^ channels, it may be assumed that spatially localized high [Ca^2+^] regions (Ca^2+^ domains) form near any individual channel when that particular channel is open (Fig. 1, left panel). In both shell and domain models, it is usually assumed that stochastic gating of L-type channels and the dynamics of the associated domains are independent except through global coupling via the bulk [Ca^2+^] and plasma membrane voltage [19]. For example, the domain model proposed and investigated by Sherman et al. [20] takes this form. Sherman et al. further assumed that Ca^2+^ domains form instantaneously when a channel activates, and collapse instantaneously when a channel deactivates or inactivates. This equilibrium formulation of domain Ca^2+^-mediated inactivation of L-type Ca^2+^ channels is viable and often utilized as an alternative to shell models. Nevertheless, when the dynamics of Ca^2+^ channel activation and inactivation are not slow compared to domain formation and collapse, the assumption of rapidly equilibrating domain [Ca^2+^] might be inadequate.

**Figure 1.**
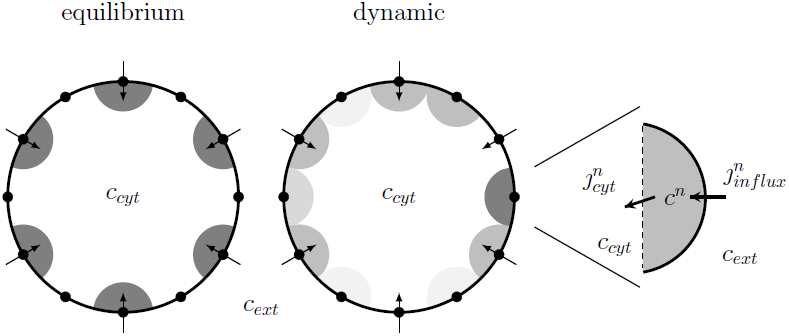
Comparison of equilibrium and dynamic domain models for Ca^2+^-mediated inactivation of L-type Ca^2+^ channels. In equilibrium domain models, low density channels are not only locally controlled, but also inactivated by a domain [Ca^2+^] that is slaved to the channel state (high concentration when open and low concentration when closed). In the dynamic domain model presented here, low density channels experience heterogeneous domain [Ca^2+^] that depend on channel state in time-dependent manner. The right panel shows the fluxes associated with a minimal formulation of single domain. Extracellular, cytosolic, and [Ca^2+^] in the *n*^*th*^ domain are denoted by *c*_*ext*_, *c*_*cyt*_, and *c*^*n*^, respectively. The domain influx rate 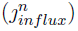 is nonzero when the Ca^2+^ channel in the *n*^*th*^ domain is open. The diffusion-mediated flux of the *n*^*th*^ domain Ca^2+^ to the cytosol is denoted by 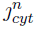

In recent years, computational models of cardiac myocytes have been developed to account for local control of Ca^2+^-induced Ca^2+^ release and heterogeneous dyadic subspace and junctional SR [Ca^2+^] [21–24]. In these models, a large number of Ca^2+^ release units (CaRUs) are simulated, each of which is represented by a discrete-state continuous time Markov chain and a compartmental representation of the dyadic subspace and junctional SR. Unfortunately, when the description of CaRU gating includes many channel states, the runtime using a Markov chain approach can be excessive.

To avoid the computationally demanding task of performing Monte Carlo simulations of a large number of CaRUs, Williams et al. [25] presented an approach to modeling local control and EC coupling in cardiac myocytes that uses probability densities to represent heterogeneous time-dependent local Ca^2+^ signals in a large number of dyadic subspaces and junctional SR domains. This approach involves numerical solution of advection-reaction equations for time-dependent bivariate probability densities of subspace and junctional SR [Ca^2+^] conditioned on CaRU state, densities that are coupled to ordinary differential equations (ODEs) for the bulk myoplasmic and network SR [Ca^2+^]. Subsequently, a moment-based approach to simulating the dynamics of local Ca^2+^ signals was found to be several orders of magnitude faster than conventional Monte Carlo simulation [26]. In this paper, we apply a population density and moment-based modeling formalism that extends the framework for domain Ca^2+^-mediated inactivation of LCCs to represent the time-dependent dynamics of domain formation and collapse (Fig. 1, middle panel). Using this modeling approach, we investigate the dependence of the inactivation function on the exponential time constant of domain collapse.

The remainder of this paper is organized as follows. First, we formulate a population density approach to modeling domain Ca^2+^-medicated inactivation of LCCs. Next, we derive the associated ODEs for the moments of these densities, and truncate and close the moment equations to produce reduced models that faithfully reproduce population density results. Using both the population density and moment-based models, we investigate the voltage-dependence of Ca^2+^-inactivation that may occur through local Ca^2+^ signaling in heterogeneous domains, and how Ca^2+^-inactivation of L-type channels may be influenced by non-equilibrium dynamics of domain formation and collapse.

## Model formulation

The compartments and fluxes included in the model formulation are shown in Fig. 1 (right panel), which includes the [Ca^2+^] in the extracellular space, the cytosol, and individual domains denoted by *c*_*ext*_, *c*_*cyt*_, and *c*^*n*^, respectively. The modeling work presented in this paper assumes that the *c*_*ext*_ and *c*_*cyt*_ are clamped. However, it is straight forward to extend the model to account for the dynamic of *c*_*cyt*_ (see Discussion). Fluxes between compartments include the influx from the extracellular space to the *n*^*th*^ individual domain 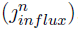, and the flux from each domain to the cytoplasm 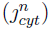.

Consistent with Fig. 1, the time-dependent dynamics of the [Ca^2+^] in the *n*^*th*^ domain is governed by the following ODE,

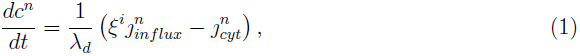

where *ξ*^*i*^ = 0 or 1 depending on whether the associated LCC is closed or open. In Eq. 1, *λ*_*d*_ = (Ω_*d*_/*β*_*d*_)/(Ω_*cyt*_/*β*_*cyt*_) is the effective volume ratio between the domain and cytoplasm that accounts for both physical volume and (constant fraction) buffering capacity. The flux from the domain to the cytoplasm is given by 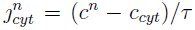. The voltage-and Ca^2+^-dependent influx, 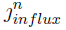, is given by Goldman-Hodgkin-Katz current equation [27].

That is, if the *n*^*th*^ LCC is open, 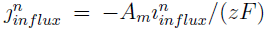 where *Am* = *C*_*m*_*β*_*cyt*_/Ω_*cyt*_ is a whole-cell capacitance scaling factor, *C*_*m*_ is the capacitive membrane area, *z* = 2 is the valence of Ca^2+^ and *F* is Faraday’s constant. The Ca^2+^ current, 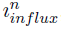, is given by 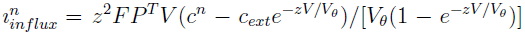 where *P*^*T*^ is the total permeability, *V* is the membrane voltage, *V*_*θ*_ = *RT*/*F*, *R* is the gas constant and *T* is the absolute temperature.

### Twelve-state LCC model

The LCC model used in this paper was introduced by Jafri et al. and reparameterized by Greenstein and Winslow [21,28]. In this model, the gating of the LCC is represented by a continuous-time, discrete-state Markov chain with twelve states, ten of which are nonconducting (closed) and two of which are conducting (open). As illustrated in Fig. 2, the upper and lower rows of states are Ca^2+^-unbound (mode normal) and Ca^2+^-bound (mode Ca), respectively. When in mode Ca, transitions to the open state *O*_*C*_*a*__ are extremely rare, because 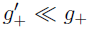. Transitions from mode normal to mode Ca depend on the rate constant *γ* = *γ*_0_*c*^*n*^, which is a linear function of the domain [Ca^2+^], that is, high [Ca^2+^] induces more transitions to mode Ca (more Ca^2+^-dependent inactivation). In both mode normal and mode Ca, there are five closed states (*C*_0_,…, *C*_4_ and *C*_*C*_*a*_0_, *C*_*C*_*a*_4_) and one open state (*O* and *O*_*C*_*a*__). Voltage-dependent transitions are determined by rate constants *α*(*V*) and *β*(*V*), which are increasing and decreasing functions of membrane voltage, respectively (see Fig. 2, caption).

**Figure 2.**
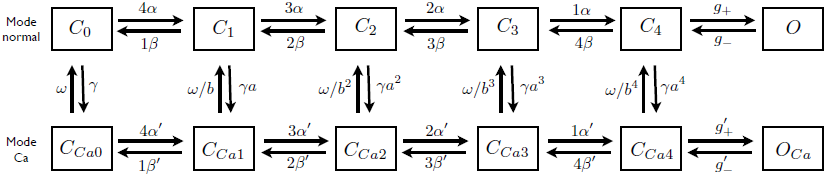
Gating scheme of the L-type channel. The 12-state L-type Ca^2+^ channel includes Ca^2+^-unbound and Ca^2+^-bound states (denoted mode normal and mode Ca, respectively). In both modes there are five closed states (*C*_0_,…, *C*_4_ and *C*_*C*_*a*_0_, *C*_*C*_*a*_4_) and one open state (*O* and *C*_*C*_*a*__). Transitions from mode normal to mode Ca depend on the rate constants *γ* (proportional to domain [Ca^2+^]) and *ω*. Voltage-dependent transitions are determined by rate constants *α*(*V*) and *β*(*V*) (mode normal) and *α′*(*V*) and *β′*(*V*) (mode Ca). Parameters follow Greenstein and Winslow [21], *α* = *α*_0_ exp(*α*_1_(*V* – *V*_0_)), *β* = *β*_0_ exp(*β*_1_(*V* – *V*_0_)), *α′* = *aα*, *β′* = *β*/*b*, *γ* = *β*_0_*c*^*n*^, = 0.85 ms^−1^, *g*_–_ = 2 ms^−1^, 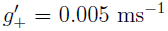, 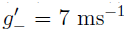, *α*_0_ = 2.0, *β*_1_ = 0.0012, *β*_0_ = 0.0882, *β*_1_ = —0.05, *a* = 2, *b* = 1.9356, *γ*_0_ = 0.44 mM^−1^ ms^−1^, *Ω* = 0.01258 ms^−1^ and *V*_0_ = 35 mV.

The transition rates between the 12 states of the LCC model can be written as a 12 × 12 infinitesimal generator matrix (*Q* matrix) that takes the form

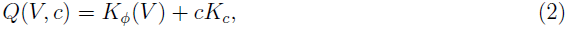

where *K*_*ϕ*_(*V*) includes the Ca^2+^-independent transitions (both voltage-dependent and voltage-independent with units of time^−1^), and *K*_*c*_ collects the association rate constants for the transitions mediated by domain Ca^2+^. Monte Carlo methods for simulating the dynamics of a population of *N* LCCs, each associated with its own domain, require numerical solution of *N* Markov chains and *N* ODEs.

### Population density formulation

We here present a population density approach to modeling the domain Ca^2+^-mediated inactivation of L-type Ca^2+^ channels that is an alternative to Monte Carlo simulation. As suming a large number (*N*) of domains, we define a continuous univariate probability density function for the domain [Ca^2+^] of a randomly sampled channel,

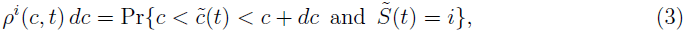

where the index *i* ∈ {*C*_0_, *C*_1_,…, *O*_*C*_*a*__} runs over the twelve states of the LCC, and the tildes on *c*̃(*t*) and *S*̃(*t*) indicate random quantities. The time-evolution of these joint probability densities is governed by the following system of advection-reaction equations [25,29–31],

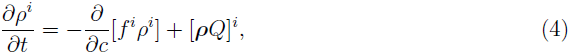

where *i* is an index over channel states, *Q* is the generator matrix given by Eq. 2, the row-vector *ρ* = (*ρ*^*C*_1_^, *ρ*^*C*_1_^,…, *ρ*^*O*_*C*_*a*__^) collects the time-dependent joint probability densities for domain Ca^2+^, and [*ρQ*]^*i*^ is the *i*^*th*^ element of the vector-matrix product *ρQ*. In Eq. 4, the reaction terms [*ρQ*]^*i*^ account for the probability flux associated with channel state changes. The advection terms of the form –*∂*(*f*^*i*^*ρ*^*i*^)/*∂c* represent the divergence of the probability flux *ϕ*^*i*^(*c*, *t*) = *f*^*i*^(*c*)*ρ*^*i*^(*c*, *t*) where the advection rate *f*^*i*^(*c*) account for the state-dependent deterministic dynamics of domain Ca^2+^,

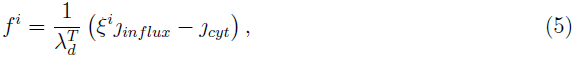

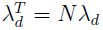 and *N* is the number of domains. As in the Monte Carlo formulation (Eq. 1), the flux term *J*_*cyt*_ in Eq. 5 is given by (*c*–*c*_*cyt*_)/*τ*. The influx term *ȷ*_*in flux*_ is linear in domain [Ca^2+^] and can be written as *ȷ*_*in flux*_ = *ȷ*_0_ – *ȷ*_1_*c* where *ȷ*_0_ = *zA*_*m*_*P*^*T*^*Vc*_*ext*_*e*^−*zV*/*V*_*θ*_^/[*V*_*θ*_(1 – *e*^−*zV*/*V*_*θ*_^)] and *ȷ*_1_ = *zA*_*m*_*P*^*T*^*Ve*^−*zV*/*V*_*θ*_^/[*V*_*θ*_(1 – *e*^−*zV*/*V*_*θ*_^)]. Consequently, the whole cell Ca^2+^ current is given by

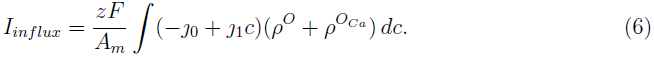

The time evolution of the joint densities *ρ*^*i*^(*c*, *t*), i.e., the dependent variables of the population density model are found by integrating Eqs. 4–5 using a total variation diminishing scheme that has been described previously [25,32]. The most important observable of the model is the probability that a randomly sampled LCC is in a given state,

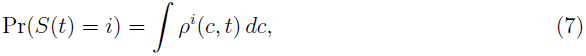

where *i* ∈ {*C*_0_, *C*_1_, …, *O*_*Ca*_}. Another important observable is the expected [Ca^2+^] in a randomly sampled domain,

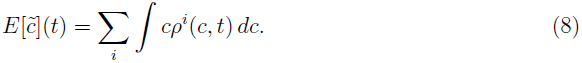

The expected [Ca^2+^] conditioned on a randomly sampled channel being in state *i* is

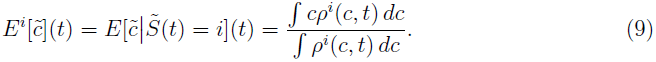

### Moment-based LCC model

The probability density approach described above is generally fast compared to Monte Carlo simulation, in part because the joint densities are univariate. However, this computational advantage diminishes when an LCC model is complex, because one joint density is required for each state. In this section, we develop a moment-based modeling approach that is computationally more efficient than the population density approach.

We begin by writing the *q*^*th*^ moment of the *i*^*th*^ joint density as

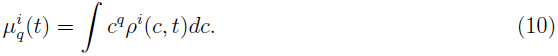

This expression implies that the zeroth moments 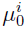 are the time-dependent probabilities that a randomly sampled channels is in state *i* (Eq. 7). The first moments, 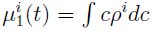 are related to the expected value of domain [Ca^2+^] conditioned on channel state though 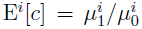 (cf. Eq. 9). The conditional variance in a randomly sampled domain is a function of the first three moments: 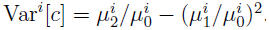.

The derivation of the moment-based LCC model begins by differentiating Eq. 10 with respect to time,

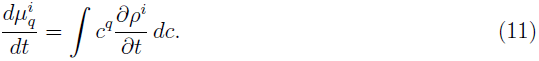

The ODEs of the moment-based model are found by replacing the factor *∂ρ*^*i*^/*∂t* in the integrand of Eq. 11 by the advection-reaction equation of the population density model (Eq. 4), which yields

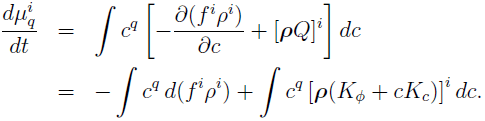

Integrating by part gives

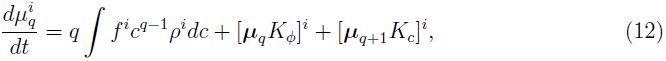

where we have eliminated boundary terms using the fact that *ϕ*^*i*^(*c*, *t*) = *f*^*i*^(*c*)*ρ*^*i*^(*c*, *t*) = 0 on the boundary (conservation of probability). We evaluate the first integral of Eq. 12 by substituting for f (Eq. 5) and simplifying,

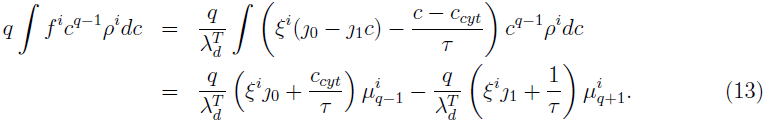

Finally, substituting Eq. 13 into Eq. 12 results in the following equation for 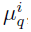,

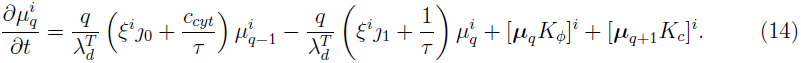

where *ξ*^*i*^ = 0 for *i* ∈ {*C*0,…, *C*4, *C*_*C*_*a*0__,…, *C*_*C*_*a*4__} and *ξ*^*i*^ = 1 for *i* ∈ {*O*, *O*_*C*_*a*__}, 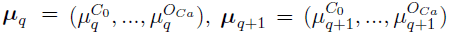, and [***μ***_*q*_ ***K***_*ϕ*_(*V*)]^*i*^ and [***μ***_*q* + 1_ ***K***_*c*_]^*i*^ are the *i*^*th*^ element of the vector-matrix product of ***μ***_*q*_ ***K***_*ϕ*_(*V*) and ***μ***_*q* + 1_ ***K***_*c*_, respectively. Note that Eq. 14 is an open system of ODEs that takes the form,

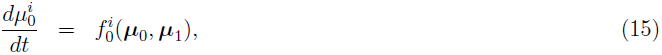

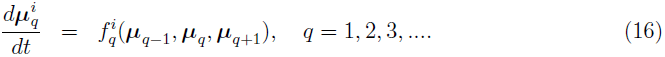

In particular, note that the equations for the *q*^*th*^ moments depend on the (*q* + 1)^*th*^ moments.

### Truncation and closure of moment ODEs

Eqs. 15–16 can be closed by assuming the (*q* + 1)^*th*^ central moment is zero, so that 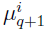 can be expressed as an algebraic function of lower moments. For example, if we assume that the conditional variance, given by 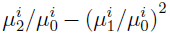, is zero for each state *i*, then the second moments are 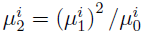. In this case, Eqs. 15–16 can be truncated and closed as follows:

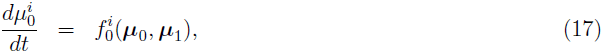

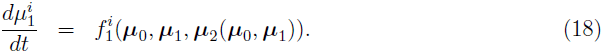

Closing the moment equations in this manner results in two ODEs per channel state—one for the zeroth moment 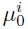, and the other one for the first moment 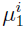 (24 ODEs in total):

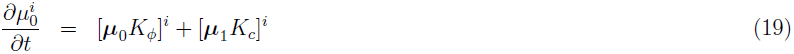

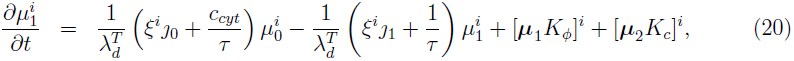

where *μ*_2_ is a row vector with elements 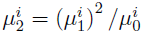.

Alternatively, we could assume the 3^*rd*^ central moments are zero. In that case, the truncated and closed moment equations take the form,

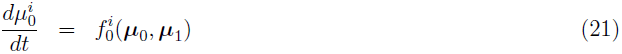

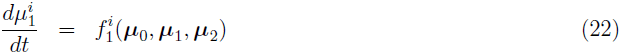

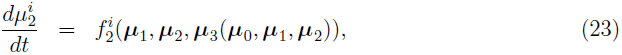

where

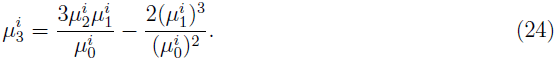

This assumption results in a moment-based model that includes 36 ODEs:

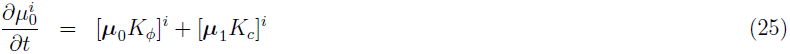

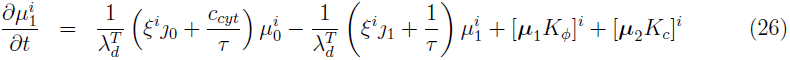

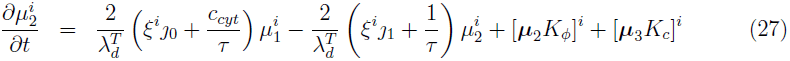

where *μ*_3_ is row vector with elements 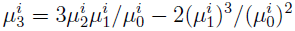.

Below, Eqs. 25–27 are referred to as the “third-order moment truncation approach” while in the sequel, Eqs. 19–20 are called the “second-order moment truncation approach.”

## Results

### Representative population density simulation results

To illustrate the population density approach to modeling domain Ca^2+^-mediated inactivation, we first show simulations of a two-pulse voltage clamp protocol, analogous to those used in the experimental quantification of Ca^2+^-inactivation of LCCs [20,33]. As shown in the top panel of Fig. 3A, the simulated command voltage began at the holding potential of *V*_*h*_ = –40 mV, and the joint densities of the model equations were equilibrated with this voltage. The command voltage was then stepped up to various prepulse potentials, *V*_*p*_, and held at *V*_*p*_ for a prescribed length of time, *t*_*p*_. The voltage was then stepped back down to the holding potential, *V*_*h*_, for duration *t*_*h*_, and then up to the test potential given by *V*_*t*_. Channel inactivation was measured by estimating the inactivation function, *h*_∝_(*V*_*p*_), defined as the normalized peak current during the test voltage pulse as a function of the prepulse potential [20],

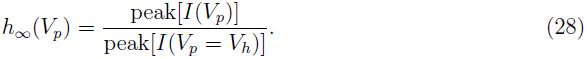

The inactivation function *h*_∝_(*V*_*p*_) gives the fraction of channels that are not inactivated and takes a value between 0 and 1. When *h*_∝_ = 1, none of the channels are inactivated; when *h*_∝_ = 0, all of the channels are inactivated.

**Figure 3.**
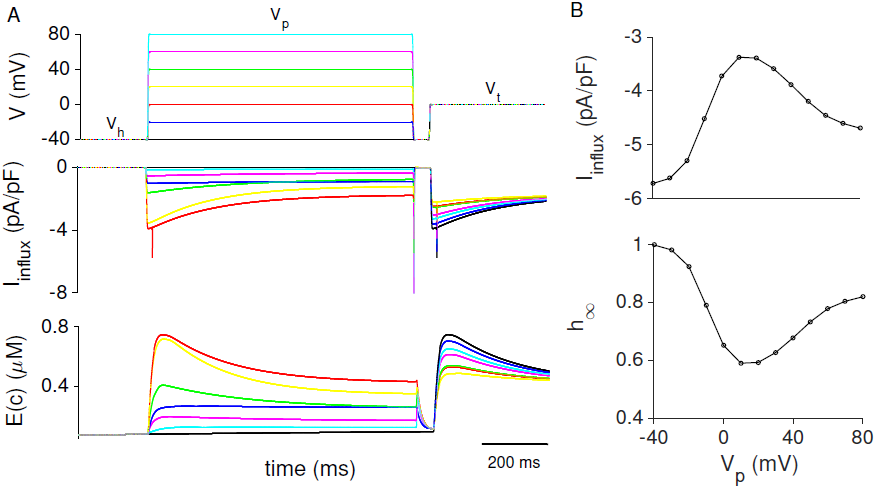
Representative simulation results. (A) The response of the whole cell current (middle panel) and expected [Ca^2+^] (bottom panel) to the two-pulse voltage clamp protocol (top panel). (B) The peak current (top panel) and the inactivation function (Eq. 28, bottom panel) to a range of prepulse potentials (–40 ≤ *V*_*p*_ ≤ 80 mV). Parameters: *V*_*h*_ = –40 mV, *V*_*t*_ = 0 mV, *V*_*p*_ = –40 to 80 mV, *τ* = 100 ms and as in Fig. 2 and Table 1.

**Table 1.** Parameters for the population density and moment-based model. See Fig. 2 for the parameters of the 12-state L-type Ca^2+^ channel.

| Symbols | Definition | Units | Value |
| --- | --- | --- | --- |
| $F$ | Faraday’s constant | coul mol<sup>-1</sup> | 96,480 |
| $R$ | gas constant | mJ mol<sup>-1</sup> K<sup>-1</sup> | 8314 |
| $T$ | absolute temperature | K | 310 |
| $V_\theta$ | $RT/F$ | mV | 26.72 |
| $P^T$ | total permeability / specific capacitance | cm<sup>3</sup> s<sup>-1</sup> $\mu\text{F}^{-1}$ | $10^{-4}$ |
| $C_m$ | capacitance | $\mu\text{F}$ | $1.534 \times 10^{-4}$ |
| $A_m$ | capacitive to volume ratio | mF L<sup>-1</sup> | 356.7 |
| $\lambda_d^T$ | effective volume ratio of domains and cytosol | - | 0.1 |
| $c_{ext}$ | extracellular $\text{Ca}^{2+}$ concentration | mM | 2 |
| $c_{cyt}$ | bulk $\text{Ca}^{2+}$ concentration | $\mu\text{M}$ | 0.1 |
| $c_{ss}$ | maximum $\text{Ca}^{2+}$ concentration (Eq. 30) | $\mu\text{M}$ | 35 |

The middle and bottom panel of Fig. 3A show the whole cell Ca^2+^ current (*I*_*in flux*_) and the mean domain [Ca^2+^] (E(*c*)) during the simulated two-pulse protocol. The largest inward currents during the test phase occurred when the prepulse voltage *V*_*p*_ was very low or very high (Fig. 3B top panel). This is consistent with the observation that during the prepulse phase little current was expressed at extreme voltages, preventing an accumulation of domain Ca^2+^ that could potentially inactivate LCCs.

The lower panel of Fig. 3B shows the inactivation curve *h*_∝_(*V*_*p*_) calculated via Eq. 28. Similar to the peak current, the inactivation function is biphasic with minimal Ca^2+^ inactivation (h ≈ 1) when the repulse potential is very low or high, and maximum Ca^2+^ inactivation (h ≈ 0.6) for intermediate repulse potentials.

Fig. 4A shows the model response to the two-pulse voltage clamp protocol using a range of domain time constants (*τ*). Slower domain time constants (large *τ*, purple line) lead to decreased inward whole cell currents during the prepulse phase (compare green and red lines). This is consistent with the observation that a slow domain time constant leads to higher expected domain [Ca^2+^] and more Ca^2+^ inactivation.

**Figure 4.**
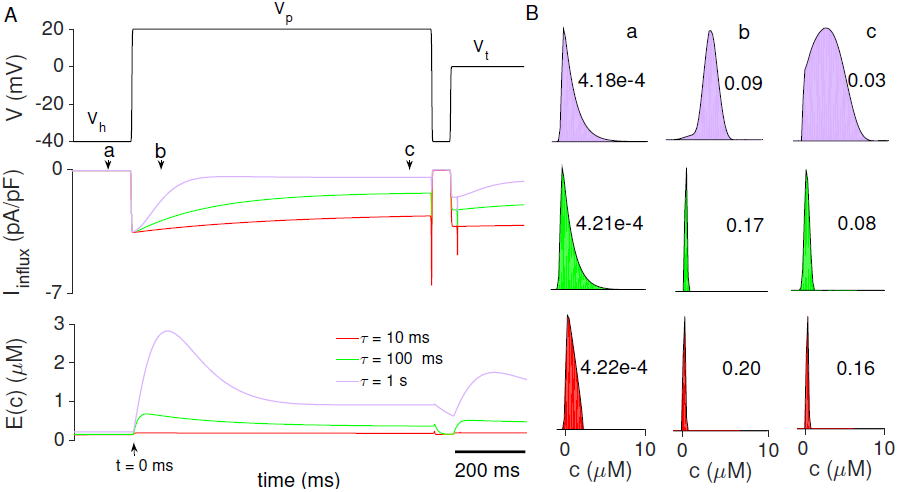
Two-pulse protocol simulation results with varied domain time constant. (A) Command voltage traces, Ca^2+^ current and expectation [Ca^2+^] when domain time constant *τ* is varied. (B) Snapshot of the sum of the joint densities for open states, *ρ*^*O*^ + *ρ*^*O*_*C*_*a*__^, at three different times (a, b, c) and three domain time constants. Parameters: *τ* = 10 ms (red), 100 ms (green) and 1 s (purple), times a, b and c are shown as arrows at –50, 80 and 750 ms, in (A), *V*_*h*_ = –40 mV, *V*_*p*_ = 20 mV, *V*_*t*_ = 0 mV and as in Fig. 2 and Table 1.

Fig. 4B shows the sum of the joint density functions of open states 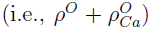 for three different domain time constants at three different times during the two-pulse protocol (arrows labeled a, b, c in panel A). Note that these densities have been normalized for clarity, so the integrated areas no longer correspond to channel open probability, which is shown as text. Consistent with Fig. 4A, the open probability at time *t* = 80 ms (b) is higher than at times *t* = –50 and 750 ms (a and c, respectively) regardless of the domain time constant. When *τ* is small (fast domain), the density functions (red and green shaded regions) are narrow and delta-function like (small variance). When *τ* is large (slow domain), the densities have greater variance (purple shaded regions).

### Comparison of population density and moment closure approaches

Fig. 5 compares the moment-based model that uses second-order and third-order truncation methods to the population density model (the correct result). When *τ* is fast or intermediate (e.g., *τ* = 100 ms), the assumption of zero variance (green) leads to nearly the same result as the population density model (+ symbols). However, when *τ* is slow (e.g., *τ* = 10 s), the result computed from the second-order moment truncation approach (khaki) deviates slightly different from the population density model (× symbols). As might be expected, this small error is eliminated using the third-order moment truncation approach (purple). Moment-based calculations in the remainder of the paper will utilize the third-order truncation method, which accurately approximate the population density model for domain time constants in the physiological range (*τ* = 1 ms to 10 s).

**Figure 5.**
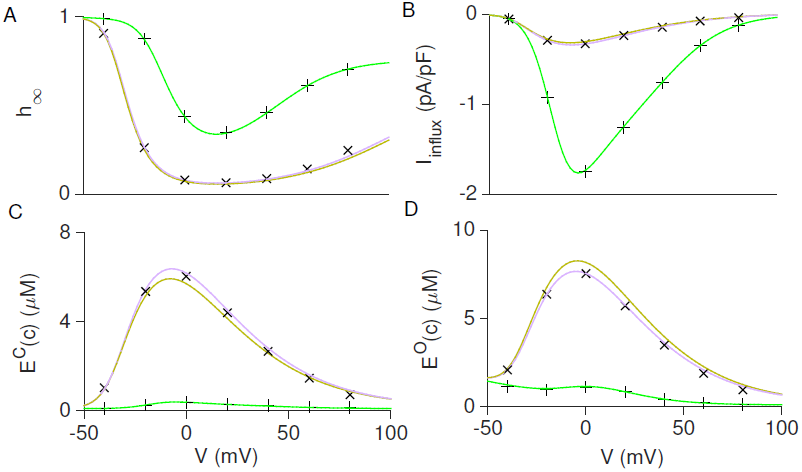
Comparison between different moment closure techniques and the population density model. Steady-state Ca^2+^-inactivation function (*h*_∝_, A), total influx current (*I*_*in flux*_, B), expected [Ca^2+^] at close state (E^*C*^(*c*), C) and open state (E^O^(*c*), D) as a function of voltage (*V*). Green and khaki lines are calculated via the second-order moment-based LCC model when *τ* = 100 ms and 10 s, respectively. Purple line is calculated via the third-order moment-based model when *τ* is 10 s. + and × symbols are computed via the population density model when *τ* is 100 ms and 10 s respectively. Other parameters as in Fig. 2 and Table 1.

### Steady-state Ca^2+^-inactivation and the domain time constant

When an LCC is open, the time-dependence of domain [Ca^2+^] can be rewritten as

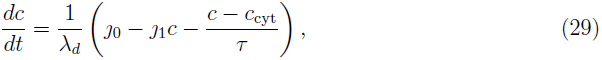

where *ȷ*_0_ and *ȷ*_1_ are defined above and we have dropped the index *n* for clarity. From Eq. 29, it is straight forward to derive the steady state domain [Ca^2+^] for and open LCC,

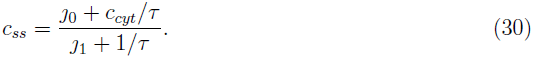

The concentration *c*_*ss*_ is the maximum [Ca^2+^] that can be achieved in a domain, its value depends on membrane voltage, the domain time constant *τ* and the total permeability *P*^*T*^, where *V* and *P*^*T*^ occur as parameters in *ȷ*_0_ and *ȷ*_1_. In this section, we investigated in how the domain time constant influences steady-state Ca^2+^-inactivation under the assumption of fixed total permeability. In the following section, we considered the related question of the domain time constant’s impact on steady-state Ca^2+^ inactivation when LCC permeability is adjusted so that the steady-state domain [Ca^2+^] (*c*_*ss*_) is fixed.

Fig. 6 shows how the domain time constant (*τ*) influences the voltage-dependence of the steady-state Ca^2+^-dependent inactivation of LCCs in the population density and moment-based models. For each domain time constant and voltage, the steady-state fraction of LCCs in four lumped states are shown, namely, mode normal open 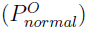, mode Ca open 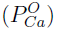, mode normal closed (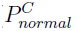, including contributions from states *C*_0_,…, *C*_4_), and mode Ca closed (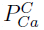, states *C*_*C*_*a*0__, …, *C*_*C*_*a*_4___). For all domain time constants studied, increasing the voltage leads to increased steady-state open probabilities 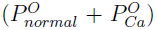. Slowing the domain time constant increases the probability that a randomly sampled channel is in mode 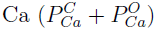 regardless of voltage, consistent with our prior observation that slower domain time constants result in higher domain [Ca^2+^] (Fig. 7C) and decreased open probability 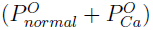.

**Figure 6.**
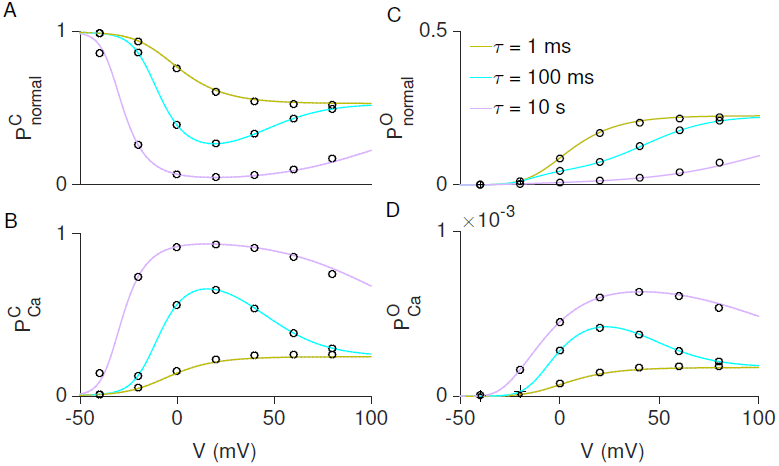
Comparison of steady-state probabilities of L-type channels states when the domain time constant *τ* is varied. The fraction of channel in closed states of mode normal 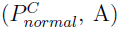 and mode 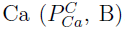, and the fraction of channels in open state of mode normal 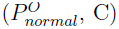 and mode 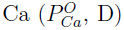, as a function of *V*_*m*_. The khaki, blue and purple lines are the simulation results of the moment-based model when *τ* = 1 ms, 100 ms and 10 s, respectively. The corresponding population density simulation results are given by open circles. Parameters as in Fig. 2 and Table 1.

**Figure 7.**
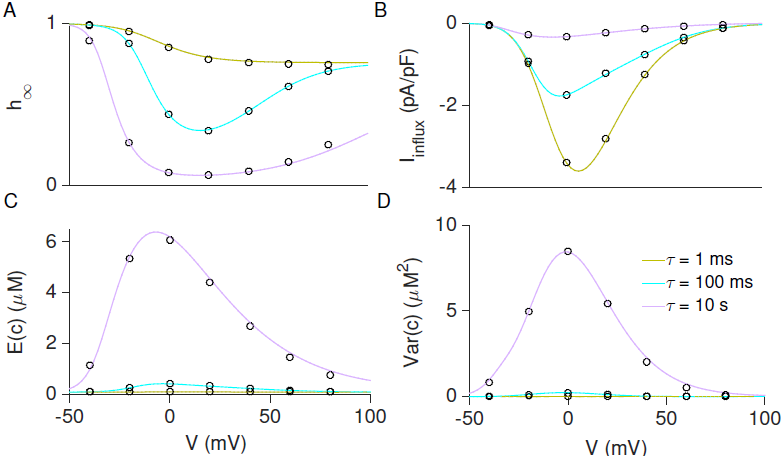
Steady-state Ca^2+^-inactivation and domain time constant *τ* with fixed *P*^*T*^. Ca^2+^-inactivation function (*h*_∝_, A), Ca^2+^ influx current (*I*_*in flux*_, B), expected [Ca^2+^] (E(*c*), C) and the variance of [Ca^2+^] in different domains (Var(*c*), D) calculated via the moment-based model as a function of *V*. The corresponding population density simulation results are given by open circles. Parameters: *τ* = 1 ms (khaki), 100 ms (blue) and 10 s (purple) and others as Fig. 2 and Table 1.

Fig. 7A shows the inactivation function (*h*_*∝*_) at steady state when τ is varied from 1 ms to 10 s. As the domain time constant *τ* increases, the inactivation function shifts downwards, corresponding to increased Ca^2+^ channel inactivation. This results from residual Ca^2+^ lingering in the domain, increasing the expected [Ca^2+^] (Fig. 7B). Although the expected domain [Ca^2+^] increases with *τ*, the total Ca^2+^ current decreases (Fig. 7B) due to decreased open probability. Fig. 7D shows that the domain Ca^2+^ concentrations are more heterogeneous (higher variance) with slow domain collapse time regardless of voltage. This is consistent with Fig. 4 where small *τ* results in narrow distribution and low variance and large *τ* yields broader distribution and higher variance.

### Ca^2^+-inactivation when maximum [Ca^2^+] is fixed

In the parameter studies of Fig. 6 and 7, the permeability *P*_*T*_ was held constant as the domain time constant *τ* was varied. Structuring the parameter study in this manner allows *τ* to influence the domain dynamics by changing the rate of domain formation and collapse as well as the steady-state domain [Ca^2+^], given by *c*_*ss*_ = (*ȷ*_0_ + *c*_*cyt*_/*τ*)/(*ȷ*_1_ + 1/*τ*). Fig. 8 presents an alternative parameter study that controls for the effect of the domain time constant on the steady state domain [Ca^2+^], thereby highlighting the manner in which the rate of domain formation and collapse influences Ca^2+^mediated inactivation of LCCs.

**Figure 8.**
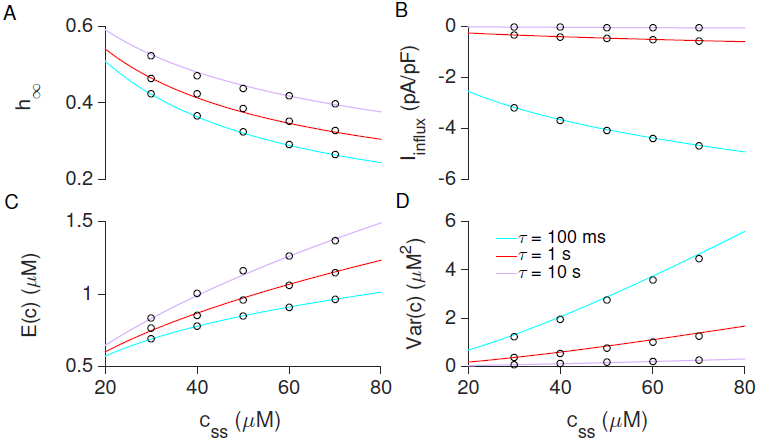
Steady-state of Ca^2+^-inactivation and domain time constant *τ* with c_ss_ and voltage fixed. *h*_∝_, *I*_*in flux*_, E(*c*) and Var(*c*) calculated via the moment-based model as a function of the maximum domain [Ca^2+^], *c*_*ss*_. The corresponding population density simulations are given by open circles. Parameters: *τ* = 100 ms (blue), 1 s (red) and 10 s (purple line), *V* = –10 mV, and others as Fig. 2 and Table 1.

Fig. 8 shows that for a given voltage and domain time constant *τ*, increasing the permeability of the channel (and thus *c*_*ss*_, the maximum domain [Ca^2+^] that can be achieved) leads to an increase in Ca^2+^-mediated inactivation (decreased *h*_∝_). On the other hand, when the permeability is adjusted so that the maximum domain [Ca^2+^] is fixed decreasing *τ* (faster domain) increases both the mean domain [Ca^2+^] (Fig. 8C) and Ca^2+^-dependent inactivation (Fig. 8A). When *c*_*ss*_ is fixed, a slower domain leads to smaller variance, i.e., Ca^2+^ channels in different domains are likely to experience similar [Ca^2+^] (Fig. 8).

## Discussion

### Summary of results

In this paper, we have shown how a population density approach (Eq. 4) to modeling Ca^2+^-mediated inactivation of L-type Ca^2+^ channels is an extension of (and improvement upon) biophysical theory that assumes that domain [Ca^2+^] is proportional to single channel current (recall Fig. 1). The population density approach is similar to traditional domain models of Ca^2+^-mediated inactivation [20] in that both assume a large number of low-density Ca^2+^ channels and a minimally represent action of the heterogeneity of domain [Ca^2+^]—a potentially important feature of Ca^2+^-mediated inactivation that is not captured by common pool models.

However, the population density approach is distinct from traditional multiscale models of Ca^2+^-inactivation in its representation of the time-dependent formation and collapse of Ca^2+^ domains associated with L-type channels. Similar to previous work focused on local control of excitation-contraction coupling in cardiac myocytes [25], the population density approach to modeling Ca^2+^ inactivation of L-type channels is often preferable to Monte Carlo simulation of the stochastic dynamics of channels and domains. This is due to the fact that the computational efficiency of a population density model scales with the number of states in the Markov chain model of the L-type channel, as opposed to the (far greater) number of channels present in the plasma membrane of the cell. Traditional equilibrium domain models also have this advantage, but do not account for the dynamics of domain formation and collapse that may in some cases influence the kinetics of Ca^2+^ inactivation [29,30].

The population density formalism allows the derivation of moment-based models of domain Ca^2+^ inactivation that are extremely computationally efficient. We have derived two different moment-based models that are distinguished by the number of ODEs per channel state retained after truncation of the open system of moment equations as well as the assumptions made to close the moment equations. Both the second-order (Eqs. 19–20, zero variance) and third-order (Eqs. 25–27, zero third central moment) moment-based models performed well when validated by comparison to corresponding population density simulations, but the third-order moment-based model was extremely accurate and valid for a wider range of domain time constants (Fig. 5). The second-order moment-based model is most accurate when the domain time constant is relatively small (fast domain, *τ* < 100 ms), because in that case the joint distributions for domain [Ca^2+^] conditioned on channel state are very focused (low variance, recall Fig. 5).

Using both the population density and moment-based models, we investigated the dependence of the steady-state inactivation of the 12-state L-type Ca^2+^ channel model [21] on the exponential time constant (*τ*) for domain formation and collapse. When the study was performed using a fixed permeability for the L-type channel, faster domains (smaller *τ*) leads to less inactivation for a wide range of clamped voltages. When the channel permeability is chosen to be a function of *τ* that results in a fixed maximum domain [Ca^2+^], a smaller domain time constant leads to increased Ca^2+^-mediated inactivation, presumably because the kinetics of domain formation subsequent to channel opening are more rapid.

### Limitations and possible extensions

Although the computational efficiency of the probability density and moment-cased calculations is notable, the runtimes of both models are proportional to the number of states in a given L-type channel model. Consequently, both methods may have little computational advantage if the LCC model of interest is extremely complex. In addition, the efficiency of the probability density approach is dependent upon the number of meshpoints used in solving the advection-reaction equations.

For simplicity, we have illustrated the population density and moment-based models under the assumption that plasma membrane fluxes do not change the bulk cytosolic [Ca^2+^] (that is, *c*_*cyt*_ is clamped). However, it is straightforward to relax this assumption and thereby allow a dynamic bulk intercellular [Ca^2+^]. For example, assuming the rate of ATP-dependent plasma membrane Ca^2+^ efflux is given by *J*_*out*_ = *k*_*out*_ *c*_*cyt*_, the ODE for bulk cytosolic Ca^2+^ is

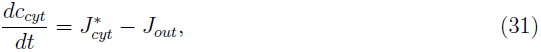

where 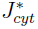 is the total flux from domains to cytosol,

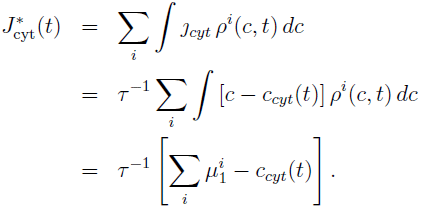

In spite of the fact that we have chosen to illustrate the population density and moment-based models through simulated voltage clamp recordings, the modeling formalism is easily modified to simulate current clamp recordings.

The population density approach presented here is well-suited to investigate whole-cell potassium currents that arise through voltage-and Ca^2+^-dependent stochastic gating of SK and BK channels, both of which play important physiological roles in the heart, brain and muscle cells and are often spatially co-localized with L-type Ca^2+^ channels [14,34–37]. Previous work by Stanley et al. [38] has shown that the stochastic gating of Ca^2+^ channels increases the activation of SK channels. Cox recently presented a Ca_v_ 2.1/BK_Ca_ model and that suggested that Ca^2+^ channels will open during a typical cortical neuron action potential, while the associated BK_Ca_ channel opens in only 30% of trials [39]. Furthermore, this percentage is sensitive to the action potential duration, the distance between the two channels in the signaling complex, and the concentration of intercellular Ca^2+^ buffers [39]. Extensions of the population density and moment-based model that account for the dynamic of Ca^2+^ buffering and the geometric relationship between channels is an important avenue for future research.

## ACKNOWLEDGMENTS

The work was supported in part by National Science Foundation Grant DMS 1121606 to GDS and the Biomathematics Initiative at The College of William & Mary.

## REFERENCES

[1] D Lipscombe, Q J Pan, and A C Gray. Functional diversity in neuronal voltage-gated Ca^2+^ channels by alternative splicing of Ca_v_*α*_1_. Mol Neurobiol, 26(1):21–44, 2002.

[2] Donald M Bers. Cardiac excitation-contraction coupling. Nature, 415(6868):198–205, Jan 2002.

[3] H Cheng, W J Lederer, and M B Cannell. Calcium sparks: elementary events underlying excitation-contraction coupling in heart muscle. Science, 262(5134):740–744, 1993.

[4] M B Cannell, H Cheng, and W J Lederer. The control of calcium release in heart muscle. Science, 268(5213):1045–1049, 1995.

[5] T H Murphy, P F Vorley, and J M Baraban. L-type voltage-sensitive calcium channels mediate synaptic activation of immediate early genes. Neuron, 7(4):625–635, 1991.

[6] J A Haack and R L Rosenberg. Calcium-dependent inactivation of L-type calcium channels in planar lipid bilayers. Biophys J, 66(4):1051–1060, 1994.

[7] T Budde, S Meuth, and H C Pape. Calcium-dependent inactivation of neuronal calcium channels. Nat Rev Neurosci, 3:873–883, 2002.

[8] D Lipscombe, T D Helton, and WF Xu. L-type calcium channels: the low down. J Neurophysiol, 92:2633–2641, 2004.

[9] L Baumann, A Gerstner, XG Zong, M Biel, and C Wahl-Schott. Functional characterization of L-type Ca^2+^ channel Ca_v_1.4a1 from mouse retina. Invest. Ophthal. Vis. Sci., 45(2):708–713, Feb 2004.

[10] E A Ertel, K P Campbell, M M Harpold, F Hofmann, Y Mori, E Perez-Reyes, A Schwartz, T P Snutch, T Tanabe, L Birnbaumer, et al. Nomenclature of voltage-gated calcium channels. Neuron, 25(3):533–535, 2000.

[11] M Simon, J F Perrier, and J Jounsgaard. Subcellular distribution of L-type Ca^2+^ channels responsible for plateau potentials in motoneurons from the lumbar spinal cord of the turtle. Eur J Neurosci, 18:258–266, 2003.

[12] A Marcantoni, P Baldelli, J M Hernandez-Guijo, V Comunanza, V Carabelli, and E Carbone. L-type calcium channels in adrenal chromatin cells: role in pace-making and secretion. Cell Calcium, 42:397–408, 2007.

[13] A Koschak, D Reimer, I Huber, M Grabner, H Glossmann, J Engel, and J Strissnig. α1D (Ca_v_ 1.3) subunits can form L-type Ca^2+^ channels activating at negative voltages. J Biol Chem, 276:22100–22106, 2001.

[14] D H Vandael, A Marcantoni, S Mahapatra, A Caro, P Ruth, A Zuccotti, M Knipper, and E Carbone. Ca_v_1.3 and BK channels for timing and regulating cell firing. Mol Neurobiol, 42:185–198, 2010.

[15] G Huang, J Y Kim, M Dehoff, Y Mizuno, K E Kamm, P F Worley, S Muallem, and W Z Zeng. Ca^2+^ signaling in microdomains: Homer1 mediates the interaction between RyR2 and Ca_v_ 1.2 to regulate excitation-contraction coupling. J Biol Chem, 282(19):14283–14290, 2014.

[16] C Christel and A Lee. Ca^2+^-dependent modulation of voltage-gated Ca^2+^ channel. Biochimica et Biophysica Acta (BBA)-General Subjects, 1820(8):1243–1252, 2012.

[17] A Torrente, P Mesirca, and P Neco. Ca_v_1.3 L-type calcium channels-mediated ryanodine receptor dependent calcium release controls heart rate. Biophys J, 100:567a, 2011.

[18] Y X Li, J Rinzel, L Vergara, and S S Stojilkovic. Spontaneous electrical and Ca^2+^ oscillations in unstimulated pituitary gonadotrophs. Biophys J, 69(3):785–795, 1995.

[19] A Zweifach and R S Lewis. Rapid inactivation of depletion-activated calcium current (ICRAC) due to local calcium feedback. J Gen Physiol, 105(2):209–226, Feb 1995.

[20] A Sherman, J Keizer, and J Rinzel. Domain model for Ca^2+^-inactivation of Ca^2+^ channels at low channel density. Biophys J, 58(4):985–995, 1990.

[21] J L Greenstein and R L Winslow. An integrative model of the cardiac ventricular myocyte incorporating local control of Ca^2+^ release. Biophys J, 83(6):2918–2945, Dec 2002.

[22] A J Tanskanen, J L Greenstein, B O’Rouke, and R L Winslow. The role of stochastic and modal gating of caridac L-type Ca^2+^ channels on early after-depolarizations. Biophys J, 88:85–95, 2005.

[23] J A Hartman, E A Sobie, and G D Smith. Calcium sparks and homeostasis in a minimal model of local and global calcium responses in quiescent ventricular myocytes. Am. J. Physiol. Heart Cir. Physiol., 299:H1996–H2008, 2010. First published Spetember 17, 2010; doi:10.1152/ajpheart.00293.2010.

[24] G S B Williams, A C Chikando, H T Tuan, E A Sobie, W J Lederer, and M S Jafri. Dynamics of calcium sparks and calcium leak in the heart. Biophys J, 101:1287–1296, 2011.

[25] G S B Williams, M A Huertas, E A Sobie, M S Jafri, and G D Smith. A probability density approach to modeling local control of Ca^2+^-induced Ca^2+^ release in cardiac myocytes. Biophys. J., 92(7):2311–2328, 2007.

[26] G S B Williams, M A Huertas, E A Sobie, M S Jafri, and G D Smith. Moment closure for local control models of Ca^2+^-induced Ca^2+^ release in cardiac myocytes. Biophys. J., 95(4):1689–1703, 2008.

[27] B Hille. Ion channels of excitable membranes. Sinauer Associates, 3rd edition, 2001.

[28] M S Jafri, J J Rice, and R L Winslow. Cardiac Ca^2+^ dynamics: the roles of ryanodine receptor adaptation and sarcoplasmic reticulum load. Biophys J, 74(3):1149–1168, 1998.

[29] R Bertram and A Sherman. Population dynamics of synaptic release sites. SIAM J Appl Math, 58(1):142–169, Feb 1998.

[30] B Mazzag, C Tignanelli, and G D Smith. The effect of residual Ca^2+^ on the stochastic gating of Ca^2+^-regulated Ca^2+^ channels. J. Theor. Biol., 235(1):121–150, 2005.

[31] M A Huertas and G D Smith. The dynamics of luminal depletion and the stochastic gating of Ca^2+^-activated Ca^2+^ channels and release sites. J. Theor. Biol., 246(2):332–354, 2007.

[32] M A Huertas and G D Smith. A multivariate population density model of the dLGN/PGN relay. J. Comput. Neurosci., 21(2):171–189, 2006.

[33] T D Plant. Properties and calcium-dependent inactivation of calcium currents in cultured mouse pancreatic B-cells. J Physiol, 404:731–747, 1988.

[34] C Vergaraa, R Latorrea, N V Marrionb, and J P Adelmanb. Calcium-activated potassium channels. Curr Opin Neurobiol., 8(3):321–329, 1988.

[35] X Y Qi, J G Diness, B J Brundel, X B Zhou, P Naud, C T Wu, J Huang, M Harada, M Aflaki, D Dobrev, M Grunnet, and S Nattel. Role of small-conductance calcium-activated potassium channels in atrial electrophysiology and fibrillation in the dog. Circulation, 129(4):430–440, 2014.

[36] R S Hammond, C T Bond, T Strassmaier, T J Ngo-Anh, J P Adelman, J Maylie, and R W Stackman. Small-conductance Ca^2+^-activated potassium channel type 2 (SK2) modulates hippocampal learning, memory, and synaptic plasticity. J Neurosci, 26(6):1844–1853, 2006.

[37] D Pribnow, T Johnson-Pais, C T Bond, J Keen, R A Johnson, A Janowsky, C Silvia, M Thayer, J Maylie, and J P Adelman. Skeletal muscle and small-conductance calcium-activated potassium channels. Muscle Nerve, 22(6):742–750, 1999.

[38] D A Stanley, B L Bardakjian, M L Spano, and W L Ditto. Stochastic amplification of Ca^2+^-activated potassium currents in Ca^2+^ microdomains. J Comput Neurosci, 31(3):647–666, 2011.

[39] D H Cox. Modeling a Ca^2+^ channel/BK_Ca_ channel complex at the single-complex level. Biophys J, 107:2797–2814, 2014.

